# Effect of Erufosine on Membrane Lipid Order in Breast Cancer Cell Models

**DOI:** 10.1101/2020.03.09.983874

**Authors:** R. Tzoneva, T. Stoyanova, A. Petrich, D. Popova, V. Uzunova, A. Momchilova, S. Chiantia

**Author notes:** Equal contribution (R. Tzoneva, T. Stoyanova, S. Chiantia).

## Abstract

Alkylphospholipids are a novel class of antineoplastic drugs showing remarkable therapeutic potential. Among them, Erufosine (EPC3) is a promising drug for the treatment of several types of tumors. While EPC3 is supposed to exert its function by interacting with lipid membranes, the exact molecular mechanisms involved are not known yet. In this work, we applied a combination of several fluorescence microscopy and analytical chemistry approaches (i.e., scanning fluorescence correlation spectroscopy, line-scan fluorescence correlation spectroscopy, generalized polarization imaging, as well as thin layer and gas chromatography) to quantify the effect of EPC3 in biophysical models of the plasma membrane, as well as in cancer cell lines. Our results indicate that EPC3 affects lipid-lipid interactions in cellular membranes by decreasing lipid packing and increasing membrane disorder and fluidity. As a consequence of these alterations in the lateral organization of lipid bilayers, the diffusive dynamics of membrane proteins are also significantly increased. Taken together, these findings suggest that the mechanism of action of EPC3 might be linked to its effects on fundamental biophysical properties of lipid membranes, as well as lipid metabolism in cancer cells.

## INTRODUCTION

Erufosine (EPC3) is a novel derivate of erucylphosphocholine that belongs to a group of antineoplastic drugs based on alkyl ether lipids (1). EPC3 can effectively be applied intravenously, can cross the blood-brain barrier and shows antitumor activity in the μM range (1, 2). For these reasons, EPC3 is a promising drug for treatment of several types of tumors, including human urinary-bladder carcinoma, breast carcinoma, glioblastoma and multiple myeloma (2). Due to its hydrophobic nature, this molecule is supposed to interact with cellular membranes, but detailed information regarding its molecular mechanism of action is scarce. On the other hand, other alkylphospholipids (APL) have been characterized more in detail. For example, it was shown that Edelfosine – one of the first characterized APLs – induces apoptosis in cancer cells via interactions with lipid rafts (1), i.e. lipid-protein domains of the plasma membrane (PM) which are enriched in sphingolipids and cholesterol (3–5) and are involved in several cellular functions (see e.g. (6, 7)). Such domains can also be characterized in protein-free model membrane systems (e.g. lipid vesicles) constituted of typical PM lipids (e.g. saturated sphingomyelin (SM), unsaturated phosphatidylcholine (PC) and cholesterol). It was shown in fact that these model membranes display a phase separation into a liquid-ordered (L_o_) and a liquid-disordered (L_d_) phase (8). The L_o_ phase is enriched in saturated lipids and cholesterol and, therefore, provides a simple physical model to study raft-like domains (9, 10). In this context, it was shown that APLs partition into lipid bilayers and directly interact with L_o_ domains (11). The observed effects vary depending on lipid composition and the investigated APL (e.g. Edelfosine or Miltefosine) and include: disorganization of L_o_ domains (11), moderate to significant increase of membrane fluidity (12, 13) and stabilization of SM/cholesterol domains (14). More specifically, studies on Erucylphosphocholine (which is more similar to EPC3, due to the shared unsaturated acyl chain structure) indicate that this APL increases the fluidity of both cellular and model membranes (12), while weakening SM-cholesterol interactions (15).

So far, studies regarding the molecular mechanisms connected to the antitumor activity of EPC3 have been limited. Recently, using spectroscopic ellipsometry as a novel technique to study solid-supported lipid model systems, we have shown that treatment of lipid films composed of PC, SM and cholesterol with EPC3 induces an increase in monolayer thickness (16).

In this work, we employed several methods based of fluorescence microscopy and analytical chemistry to investigate for the first time how EPC3 affect the physical properties of the PM. First, we have applied line-scan fluorescence correlation spectroscopy (lsFCS) (17, 18) and scanning fluorescence correlation spectroscopy (sFCS) (19–21) to monitor the dynamics of membrane lipids and proteins, both in simple artificial membranes and directly in the PM of living cells. These methods belong to the family of fluorescence fluctuation techniques and were effectively used in the past to quantify membrane dynamics (and, indirectly, membrane order) (22). Furthermore, the use of supported lipid bilayers (SLBs) as membrane models allows the study of lipid-lipid interactions in a controlled environment and in specific relation to the precise lipid composition of the bilayer (23, 24).

Second, using thin layer chromatography and gas chromatography, we evaluated the changes occurring in phospholipid and cholesterol amounts in the membranes of EPC3-treated cancer cells.

Finally, we quantified the influence of EPC3 on membrane order for the PM of cancer cell models by using polarity-sensitive fluorescent probes (25). More in detail, we have investigated the spectral properties of two membrane probes (Laurdan and Di-4-ANEPPDHQ) which were shown to be influenced by lipid packing, membrane hydration and composition to different extents (25, 26).

Our results indicate that EPC3 modulates the PM lipid composition of cancer cells. Furthermore, it affects lipid-lipid interactions both in lipid membrane models and cellular membranes. Such alterations in membrane order appear to have a direct effect on membrane protein dynamics.

## MATERIALS AND METHODS

### Chemicals

Erufosine (EPC3) was synthesized in the Max Planck Institute for Biophysical Chemistry, Göttingen, Germany, and was most graciously provided by Prof. Martin R. Berger. It was dissolved in a PBS (phosphate buffer saline, pH 7.4) in 10 mM stock solution and kept at 4°C. Di-4-ANEPPDHQ and Laurdan (6-dodecanyl-2-dimethylaminonaphtalene) were from Molecular Probes (Eugene, OR). L-α-phosphatidylcholine from chicken egg (EggPC), cholesterol from ovine wool (Chol), sphingomyelin from porcine brain (bSM), 23-(dipyrrometheneboron difluoride)-24-norcholesterol (TF-Chol) and 1,2-dioleoyl-sn-glycero-3-phosphoethanolamine-N-(lissamine rhodamine B sulfonyl) (Rhod–DOPE) were purchased from Avanti Polar Lipids, Inc. (Alabaster, AL).

### Cell culture

MDA-MB-231 highly invasive breast cancer cells and MCF-7 epithelial cancer cells were acquired from the American Type Culture Collection (ATCC, USA). Both cell lines are derived from breast adenocarcinoma, with MCF-7 cells retaining some characteristics of the differentiated mammary epithelium.All cell lines were incubated in phenol red-free DMEM culture medium with 10% fetal bovine serum, 2 mM L-Glutamine, and 100 U/mL penicillin and 100 μg/mL streptomycin at 37 ◦C and 5% CO2. Cells were passaged every 3–5 days, no more than 15 times. All solutions, buffers and media used for cell culture were purchased from PAN-Biotech (Aidenbach, Germany).

### Plasmids

The plasmid coding for human EGFR tagged with EGFP (hEGFR-EGFP) was a kind gift from Alexander Sorkin (Addgene plasmid #32751) (27). To replace EGFP with mEGFP, both plasmids hEGFR-EGFP and mEGFP-N1 (gift from Michael Davidson, Addgene plasmid #54767) were digested with AgeI and NotI (New England Biolabs GmbH, Ipswich, MA). The hemagglutinin (HA) gene from influenza virus strain A/FPV/Rostock/34 (H7N1) tagged with mEGFP at the C-terminus (FPV-HA-mEGFP) was cloned based on the previously described FPV-HA-mEYFP (gift from Andreas Herrmann, Humboldt University Berlin) (28). Briefly, FPV-HA-mEYFP was digested using BglII and SacII (New England Biolabs GmbH, Ipswich, MA), and the obtained HA-insert was ligated into mEGFP-N1. This plasmid is available on Addgene (#127810).

### Supported lipid bilayers

Supported lipid bilayers (SLBs) were prepared as previously described (29, 30). Briefly, lipids were mixed in organic solvent at the desired concentrations and dried on the walls of a glass vial. The lipid film was then rehydrated in PBS pH 7.4 and, after vigorous vortexing, sonicated to clarity in a bath sonicator. Typical concentrations during sonication were ⁓5-10μM. The thus obtained small unilamellar vesicle suspension was then diluted ca. 10-fold and 100 μL were deposited on a small thin piece (⁓10 mm^2^) of freshly-cleaved mica glued to the surface of a glass coverslip (thickness #1). The mica and the vesicles suspension were confined using a 7 mm-plastic cylinder, also glued to the glass surface. Vesicle fusion and bilayer formation were induced by addition of 3 mM CaCl_2_. The volume was adjusted to 300 μL and the suspension was incubated for 10 min. Unfused vesicles were removed by addition and removal of 500 μL DPBS, performed 10 times. The treatment with EPC3 was performed by adding the drug at the desired concentration in PBS directly on top of the SLB and waiting ca. 30 min before performing fluorescence measurements.

### Line-scan Fluorescence Correlation Spectroscopy on SLBs

Line-scan FCS (lsFCS) measurements on SLBs were performed as previously described (17). Briefly, data were acquired by scanning repeatedly the focal volume in a linear fashion in the plane of the membrane. Line scans of ca. 5-10 μm length were chosen so that both bilayer phases (L_d_ and L_o_) were scanned through. We typically acquired 250000 lines, each divided in 256 pixels, with a pixel time of 1.27 μs (line time: 763 ms). Intensity values were correlated along each line and between different lines, calculating the full spatio-temporal autocorrelation G(ξ, τ_i_). To account for photobleaching, a mathematical correction was applied (17). Data analysis was performed with a custom-written script in Matlab, by fitting G(ξ, τ_i_) using a weighted nonlinear least-squares fitting algorithm and a mathematical model taking into account the linear scanning and 2-d Brownian diffusion. To capture the statistical information at larger lag times, we also included in the evaluation (via global fitting) the analysis of the temporal autocorrelation curve G(0, τ_i_), calculated on a logarithmic scale with a multiple τ-algorithm (17). We could thus obtain estimates for the waist ω_0_ and the diffusion coefficient D for a fluorescent lipid analogue (TF-Chol, 0.01 mol%) added to the examined SLBs. Autocorrelation curves were calculated independently for the different lipid phases, which could be identified due to the distinct affinity of a fluorescent dye (i.e. Rhod-DOPE, 0.1 mol%) to the L_o_- and the L_d_-phases. Line scans were performed using the same setup described above for sFCS, using a 488 nm argon laser (ca. 1.5 μW) for the excitation of TF-Chol. The signal originating from Rhod-DOPE (excitation: 561 nm, 561/488 dichroic mirror, emission collected between 571 and 650 nm) was collected only in order to distinguish the lipid phases, but not further used for lsFCS analysis. The D values for TF-Chol in the L_o_ and L_d_ phase of SLB, in the absence of EPC3, were calculated as average of three independent measurements performed in different days. These values were then used as normalization reference for D values measured in the presence of EPC3, so to emphasize the effect of the drug rather than day-to-day variations.

### Lipid extraction and analysis of phospholipids

MCF-7 and MDA-MB-231 cells were seeded in 25-cm^2^ cell culture flasks at a density of 1,5×10^5^ cells/ml. After a 24 h incubation, cells were treated with IC50 and IC75 amounts of EPC3 (31) and further incubated for 24 hours. The extraction of membrane lipids was performed as described previously (32) with chloroform/methanol, according to the method of Bligh and Dyer (33). Briefly, the organic phase obtained after extraction was concentrated and analyzed by thin layer chromatography. The individual phospholipid fractions were separated on silica gel G 60 plates (20×20 cm, Merck, Germany) in a solvent system containing chloroform/methanol/acetic acid/d.H2O (70:35:8:4, v/v). The location of the separated fractions was visualized by iodine staining. The spots were scraped and quantified by estimation of inorganic phosphorus (34).

### Determination of cholesterol by gas chromatography

Cholesterol content in the membranes of EPC3-treated breast cancer cells (as described above) was determined using gas chromatography (35) using Carlo Erba gas-chromatograph equipped with a flame-ionization detector isothermally at 190°C and with a 2m column coated with 10% DEGS on Chromosorb W60-80mesh (Pharmacia), with nitrogen as the gas-carrier.

### Confocal microscopy imaging using di-4-ANEPPDQ and Laurdan

The cells were seeded in 35-mm microscopy dishes (CellVis, Mountain View, CA) with an optical glass bottom (#1.5 glass, 0.16 - 0.19 mm) at a density of 5×10^4^ cells/well for MDA-MB 231 and 1×10^5^ cells/well for MCF-7. After a ⁓12 h incubation, MDA-MB-231 cells were treated with a 20 (or 30) μM EPC3 solution in culture medium, corresponding to the previously measured IC50 and IC75 values (31). MCF-7 cells were treated with a 40 (or 60) μM EPC3 solution (corresponding to IC50 and IC75 values, Tzoneva R., unpublished data) in culture medium. In both cases, the treatment lasted for 24, 48 or 72 h. Control cells were incubated just with culture medium, following the same protocols. After the incubation period, the cell medium was removed. The cells were then washed twice with PBS, pH 7.4. Afterwards, 2 ml of serum-free and phenol red-free DMEM and fluorescent dye (with final concentration of 1 μM for di-4-ANEPPHQ or 5 μM for Laurdan) were added to the cell dish. The cells were further incubated for 30 min at 37 °C.

A Zeiss LSM 780 system (Carl Zeiss, Oberkochen, Germany) was used to acquire the confocal images, with a pixel size of ca. 200 nm (512 × 512 pixels). Samples were imaged using a Plan-Apochromat 40x/1.2 Korr DIC M27 water immersion objective. The excitation sources were a 488 nm argon laser (for di-4-ANEPPDQ) or a 405 nm diode laser (for Laurdan). Fluorescence was detected between 498-579 nm (Channel 1) and 620-750 nm (Channel 2) for di-4-ANEPPDQ, after passing through a 488 nm dichroic mirror, using a gallium arsenide phosphide (GaAsP) detector. Fluorescence of Laurdan was detected in the spectral ranges 410-463 nm (Channel 1) and 472-543 nm (Channel 2), after passing through a 405/565 nm dichroic mirror. Out-of-plane fluorescence was reduced by using a 42.4-μm pinhole in front of the detector.

Confocal images were analyzed as previously described (25, 26). Briefly, for each experimental condition, ca. 5-10 confocal images of treated cells were acquired. In each image, several regions of interest (ROIs) were selected in correspondence of the plasma membrane of different cells. Experiments were performed as independent duplicates on different days so that, for each EPC3 concentration and each time point, at least 50 cells were analyzed in total. All the pixels from ROIs collected within equivalent samples (i.e. same EPC3 concentration and same time point) were pooled together. For each pixel, the generalized polarization (GP) value was calculated as defined in Refs. (26), using a custom-written Matlab (The MathWorks, Natick, MA) script and setting the calibration factor G=1. The obtained results are shown as normalized occurrence histograms of all selected pixels.

### Scanning Fluorescence Correlation Spectroscopy (sFCS) in living cells

For one-color sFCS experiments, 8 × 10^4^ MDA-MB-231 cells were seeded in 35-mm dishes (CellVis, Mountain View, CA) with optical glass bottoms (#1.5 thickness, 0.16 - 0.19 mm). After 24 h, cells were treated with medium containing 30 μM EPC3. After an additional incubation for 24 h, cells were transfected by using 200 ng (mp-mEGFP and mp-mEGFP(2x)) or 600 ng (hEGFR-EGFP or A/FPV-HA-mEGFP) plasmid DNA per dish with Lipofectamine™ 3000 according to the manufacturer’s instructions (Thermo Fisher Scientific). Plasmids were incubated for 15 min with 2 μl P3000 per μg plasmid and 2 μl Lipofectamine™ 3000 diluted in 50 μl serum-free medium, and then added dropwise to the cells.

After 24 h incubation, scanning fluorescence correlation spectroscopy (sFCS) measurements were carried out on a Zeiss LSM 780 system, equipped with a Plan-Apochromat 40x/1.2 Korr DIC M27 water immersion objective. Samples were excited with a 488 nm argon laser and the fluorescence was detected between 499 and 597 nm, after passing through a 488 nm dichroic mirror, using a GaAsP detector. To decrease out-of-focus light, a pinhole size of one airy unit (~ 39 μm) was used. To keep photobleaching below 20 %, a laser power of 1.2 μW was chosen. For cell measurements, a line-scan of 256×1 pixel (pixel size 80 nm) was performed perpendicular to (i.e. across) the plasma membrane with a 403.20 μs scan time (0.67 μs pixel dwell time). For each measurement, 400000 lines were acquired in photon counting mode, and the total scan time was around 3 min per measurement. Scanning data were exported as TIFF files. At the beginning of each measurement day, the signal was optimized by adjusting the collar ring of the objective to the maximal count rate for an Alexa Fluor® 488 (AF488, Thermo Fischer, Waltham, MA) solution (50 μM dissolved in water) excited at the same laser power. For the focal volume calibration, a series of point FCS measurements was performed (ten measurements at different locations, each consisting of 15 repetitions of 10 s), and the data were fitted with a three-dimensional model including a triplet contribution. The structure parameter was typically around 6 to 9, and the diffusion time around 35 to 40 μs. The waist ω_0_ was calculated from the measured average diffusion time (τ_d,AF488_) and previously determined diffusion coefficient of the used dye at room temperature (D_AF488_ = 435 μm^2^s^−1^) (36), according to the following equation:

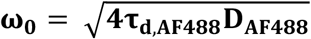

Typical values were 200-250 nm. All measurements were performed at room temperature.

sFCS analysis follows the procedure described previously (19, 21, 37). Briefly, the TIFF files were imported and analyzed in Matlab using a custom-written code. All scanning lines were aligned as kymographs and divided in blocks of 1000 lines. In each block, lines were summed up column-wise and the x position with maximum fluorescence was determined, by fitting with a Gaussian function. This algorithm finds the position of the plasma membrane in each block and is used to align all lines to a common origin. The pixels corresponding to the membrane were defined as pixels which are within ±2.5σ of the peak. In each line, these pixels were integrated, providing the membrane fluorescence time series F(t). A background correction was applied by subtracting the average pixel fluorescence value on the inner side of the membrane multiplied by 2.5σ (in pixel units) from the membrane fluorescence, in blocks of 1000 lines (38). In order to correct for depletion due to photobleaching, the fluorescence time series was fitted with a two-component exponential function and a mathematical correction was applied (17). Finally, the normalized ACF was calculated as:

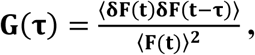

where

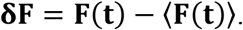

To avoid artefacts that can be caused by long-term instabilities of the system or single bright events, correlation functions were first calculated segment-wise (10 segments per time trace), and segments with distortions were manually removed before averaging the correlation functions. Eventually, a model for two-dimensional diffusion in the membrane and a Gaussian focal volume geometry (37) was fitted to the ACF:

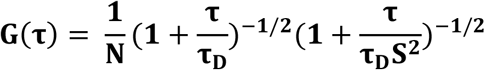

where τ_D_ denotes the diffusion time, and N the number of particles. The structure parameter S was fixed to the value of the daily based calibration measurement. Diffusion coefficients (D) were calculated using the calibrated waist ω_0_ of the focal volume: 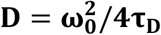.

All resulting data were analyzed using GraphPad Prism 5.0 (GraphPad Software Inc., San Diego, CA), and displayed as box plots indicating the median values and with Tukey whiskers. Quantities in the main text are expressed as mean ± SD. Sample sizes and p-values are indicated in figure captions. Statistically significant differences between control and test samples were determined using the one-way ANOVA analysis followed by the Bonferroni’s multiple comparisons test. A p-value <0.01 was considered indicative of statistical significance.

## RESULTS AND DISCUSSION

### EPC3 increases membrane fluidity in lipid bilayer models

In order to assess the influence of EPC3 on lipid-lipid interactions, we have first investigated its effects on controlled membrane models. More in detail, we have prepared SLBs mimicking the general composition of the outer leaflet of the PM. These bilayers were composed of a natural mixture of PC (i.e. EggPC), sphingomyelin (bSM) and cholesterol (4/4/2 molar ratio). Such ternary lipids mixtures are known to separate into a liquid disordered (L_d_) and a liquid-ordered phase (L_o_), thus providing a simple model for the study of phase separation occurring at the PM of living cells (22, 39, 40).

In this experiment, we first observed the effect of EPC3 on the general appearance of the L_o_-L_d_ phase separation in SLBs, via confocal fluorescence microscopy. For this purpose, SLBs were labelled with an unsaturated fluorescent lipid (Rhod-DOPE) which readily partitions in the L_d_ phase (41). As apparent in Fig. 1 A, SLBs showed bright patches enriched in Rhod-DOPE (i.e. L_d_ phase) and dark regions devoid of Rhod-DOPE (i.e. L_o_ phase). In the presence of increasing amounts of EPC3 in solution (Fig. 1 B-D), the surface occupied by the L_d_ phase increases. At the highest EPC3 concentration (i.e. 10 μM, Fig.1 D), only small patches of L_o_ phase are still visible. These results indicate that EPC3 can effectively insert into the bilayer and destabilize the L_o_ phase. A similar induction of lipid mixing was observed also for other APLs in giant unilamellar vesicles (42).

**Figure 1:**
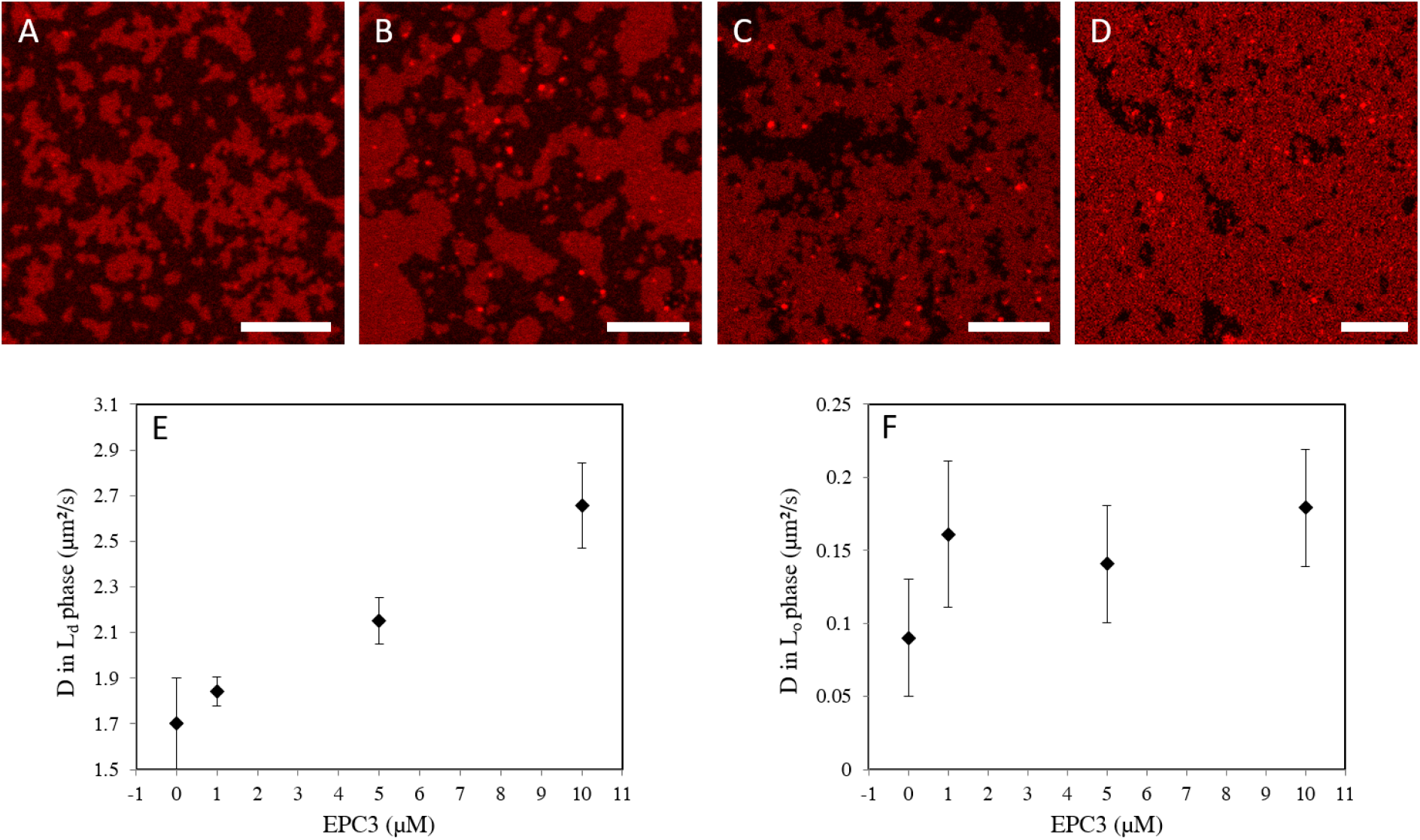
Effect of EPC3 on phase-separating model membranes. Panels A-D show representative confocal fluorescence microscopy images of EggPC/bSM/Chol 4/4/2 molar ratio SLBs in the presence of 0, 1, 5 and 10 μM EPC3, respectively. The bilayers were labelled with 0.1 mol% Rhod-DOPE. Darker zones (devoid of Rhod-DOPE) indicate L_o_ regions of the SLB. Images were acquired at RT. Scale bars are 10 μm. Panels E and F show the values of the diffusion coefficients D measured via lsFCS in the L_d_ (E) and L_o_ phase (F) of EggPC/bSM/Chol 4/4/2 molar ratio SLBs in the presence of EPC3. The reported values refer to the diffusion of a fluorescent cholesterol analogue (TF-Chol, 0.01 mol%). Independent experiments were repeated in triplicates and averaged after normalization (see Materials and Methods). Error bars represent standard deviations.

Next, we characterized the degree of order of the lipid bilayer, in each phase. To this aim, we quantified the diffusion coefficient (D) of a fluorescent membrane probe (TF-Chol) both in the L_o_ and L_d_ phases. Lipid dynamics are in general connected with membrane order, with lower D values being associated to tighter lipid packing and stronger lipid-lipid interactions (22). Using lsFCS, we measured D for TF-Chol in the L_d_ (Fig. 1 E) and L_o_ phases (Fig.1 F), in the presence of increasing concentrations of EPC3. In the absence of the drug, D was ca. 0.1 μm²/s in the ordered phase and ca. 20-fold higher in the disordered phase, as expected from previous experiments (39). In the presence of EPC3, we observed a ca. 2-fold increase in lipid diffusion dynamics in the L_d_ phase. For the L_o_ phase, we observed a similar behavior, although the total change in D was minor compared to the data spread, in this case. These results clearly suggest that EPC3 has a fluidizing effect on the lipid bilayer that leads to a significant increase of the diffusion of membrane components. On the other hand, it is not possible to determine from these data whether EPC3 interacts preferentially with (and inserts in the bilayer through) the L_o_ or the L_d_ phase. Of interest, saturated alkylphospholipids were suggested to partition at the boundary between L_o_ and L_d_ phases (42).

### EPC3 treatment alters phospholipid and cholesterol contents in cell membranes

To monitor the influence of EPC3 on the concentrations of the major phospholipid components and cholesterol in cell membranes, we characterized two breast cancer cell models: the high-invasive MDA-MB-231 cell line and the low-invasive MCF-7 cell line. Both cell types were treated with EPC3 (IC50 and IC75) for 24h and lipid amounts were measured by means of TLC and gas chromatography.

Some APLs are known to inhibit the synthesis of key phospholipids such as PC and SM (43–46). Accordingly, we observed a similar effect in EPC3-treated MCF-7 and MDA-MB-231 cells (Fig. 2 A and C). The SM content significantly decreased with increasing concentration of EPC3 in both cell lines. For instance, the decrease of SM content after treatment with IC75 EPC3 was ⁓24 % for MCF-7 cells and ⁓16% for MDA-MB-231 cells, compared to control samples. This finding is in line with the observations of Marco and coworkers (47) showing that treatment of human hepatoma HepG2 cell line with Miltefosine leads to inhibition of SM metabolism. Disturbances in SM synthesis might cause the accumulation of ceramide and sphingosine, which regulate cellular functions such as proliferation, gene expression, differentiation, mitosis, cell survival and apoptosis. For example, sphingosine and ceramide can induce apoptosis via the intrinsic apoptotic pathway by modulating the permeability of the mitochondrial membrane, thereby releasing proteins such as cytochrome C (48).

**Figure 2:**
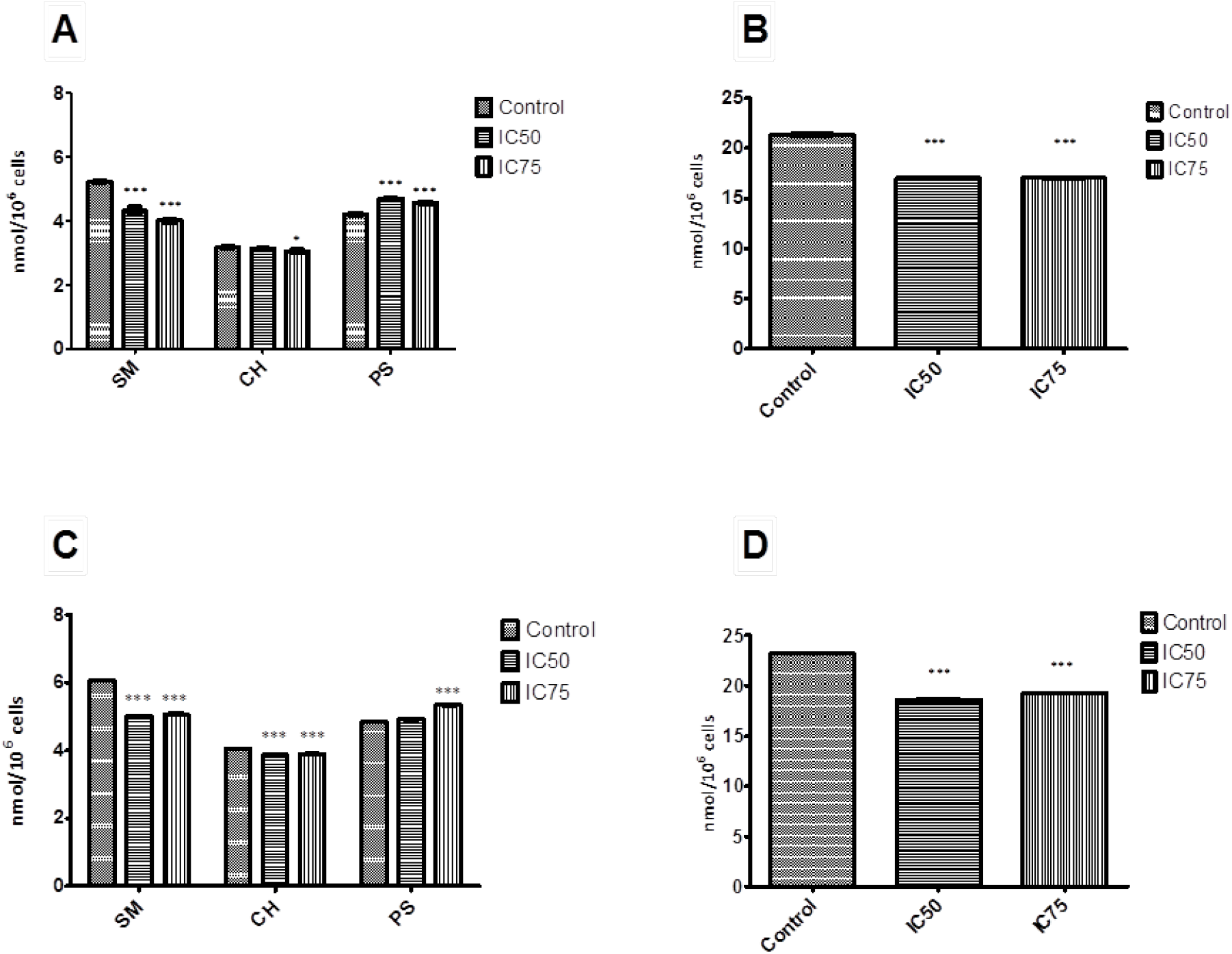
Effect of EPC3 on the level of sphingomyelin, cholesterol, phosphatidylserine and phosphatidylcholine in cell membranes. A) Amounts of sphingomyelin (SM), cholesterol (CH) and phosphatidylserine (PS) in MCF-7 cells treated with EPC3 for 24 h. B) Amounts of phosphatidylcholine (PC) in MCF-7 cells treated with EPC3 for 24 h. C) Amounts of sphingomyelin (SM), cholesterol (CH), phosphatidylserine (PS) in MDA-MB-231 cells treated with EPC3 for 24 h. D) Amounts of phosphatidylcholine (PC) in MDA-MB-231 cells treated with EPC3 for 24 h. Data are obtained as means of three independent experiments. Error bars represent the standard deviations. Statistical significance is calculated against controls in each group using ANOVA one-way test and Bonferroni post-test. *- p<0.01, ***- p<0.0001.

Furthermore, our results show a EPC3-induced reduction of PC amount. The effect was observed in both cancer cell lines (Fig. 2 B and C). In MCF-7 cells treated with IC75 EPC3, the PC decrease was 20%, compared to control samples. In MDA-MB-231 cells we observed a 17% decrease. Of interest, one of the pathways for inducing apoptosis in cells treated with APLs is thought to be indeed the inhibition of PC synthesis. Our results support this hypothesis, as they show statistically significant reductions in PC levels at both EPC3 concentrations. Previous studies have demonstrated that Miltefosine also inhibits PC synthesis (44).

In contrast, PS levels in the membranes of both treated cell lines appear to slight increase (Fig. 2 A and C). Increased PS levels have also been previously observed in cancer cells as response to chemotherapy or radiation treatment (49).

As shown in Fig. 2 A and C, treatment with EPC3 also resulted in a slight decrease in cholesterol levels. The membranes of the highly invasive cell line MDA-MB-231 were found to initially contain more cholesterol than those of MCF-7 cells. These results are consistent with the hypothesis that high cholesterol levels in cell membranes can enhance cell migration in cancer models, including MDA-MB-231 cells (50–52). Our data further show a reduction in cholesterol levels after treatment with EPC3 cells that is correlated with reduced cell survival, as we have previously observed (31). Recent studies indicate that the administration of membrane-active APLs such as Edelfosine, Erucylphosphocholine and Perifosine reduces the proliferation of HepG2 cells, disrupting cholesterol trafficking from the PM to intracellular membranes and decreasing the esterification of cholesterol (53). Our data show that treatment of both MCF-7 and MDA-MB-231 cells with highly cytotoxic concentrations of EPC3 (IC75, (31)) resulted in a small but reproducible reduction in cholesterol levels (i.e. ~5%). The molar ratio of phospholipid-to-cholesterol content of low invasive untreated MCF-7 cells was ~9.7. After treatment with EPC3 (IC75), the ratio decreased to ~8.4 (unpaired t-test, p<0.0004). The high-metastatic cell line MDA-MB-231 showed an average phospholipid-to-cholesterol ratio of ~8.4, which decreased to 7.6 upon treatment with IC75 EPC3 (unpaired t-test, p<0.0002). Since the relative amounts of cholesterol and saturated sphingolipids (e.g. SM) are fundamental determinants of the physical properties of ordered PM domains (e.g. raft domains) (54), we focused on these two membrane components. In both breast cell lines, a decrease in SM/cholesterol ratio was observed. For instance, in non-treated MCF-7 cells, this ratio was ~1.6. After treatment with IC75 EPC3, the ratio decreased to ~1.3 (unpaired t-test, p<0.0002). Untreated MDA-MB-231 cells showed a SM/cholesterol molar ratio of ~1.5. After treatment with IC50 or IC75 EPC3, the ratio decreased to ~1.3 (unpaired t-test, p<0.0001).

Altered levels of (intra-)cellular lipids and cholesterol are linked to cancer aggressiveness (55–57). More specifically, saturated lipids were suggested to reduce the fluidity and dynamics of the membrane and increases resistance to conventional chemotherapy (58). Also, reducing cholesterol content with membrane-depleting agents or inhibitors of cholesterol synthesis (e.g. statins) was suggested to alter the structure of lipid rafts (58). The consequent raft destabilization might, in turn, interfere with the proliferation and migration of tumor cells (59, 60). EPC3 treatment of both cancer cell lines induces a significant decrease in the amounts of specific cellular lipids – such as SM and cholesterol – as well as alterations in their relative molar ratios. We argue therefore that the cytotoxic effects of EPC3 might be linked to alterations in the lateral organization of cellular membranes and, in particular, to a decreased stability and/or amount of ordered lipid domains.

### EPC3 alters the fluidity of the plasma membrane of living cells

In order to investigate the effects of EPC3 directly in the PM of living cells, we applied an approach based on the spectroscopic properties of two fluorescent molecules (i.e. Di-4-ANEPPDHQ and Laurdan) which are strongly sensitive to membrane order and lipid packing (25, 26). In the context of studies regarding the effect of APLs on lipid bilayers, Laurdan was used to detect and increase in the fluidity of model membranes induced by Miltefosine (61).

In this experiment, we characterized directly the PM of both MDA-MB-231 and MCF-7 breast cancer cell models. In previous studies, we observed that EPC3 causes increased cytotoxicity, apoptosis and cytoskeleton reorganization especially in highly invasive MDA-MB-231 cells, compared to MCF-7 samples (62, 63). In addition, MDA-MB-231 cells showed cell cycle arrest after treatment with EPC3 (63).

The mechanism by which Laurdan and Di-4-ANEPPDHQ detect changes in the local membrane environment is similar for the two molecules: the less polar environment of the ordered bilayer phase (e.g. L_o_ phase or a raft-like domain in the PM) induces a blue shift in the emission maxima of both fluorescent probes. This shift can be quantified by calculating a ratiometric measurement of the fluorescence intensity observed in two spectral regions (or channels), known as a GP value. Higher GP values correspond to a relatively higher fluorescence emission in the shorter-wavelength spectral region and, therefore, a higher degree of membrane order and tighter lipid packing (26).

Fig. 3 shows normalized histograms for Di-4-ANEPPDQH GP values measured in MDA-MB-231 cells, in the presence of 20 μM or 30 μM EPC3. Cells were observed via confocal fluorescence microscopy after 24 h, 48 h and 72 h (see Fig. S1). In all cases, the PM region of several cells was manually selected and the GP values for each pixel within these ROIs were calculated as described in the Materials and Methods section.

**Figure 3:**
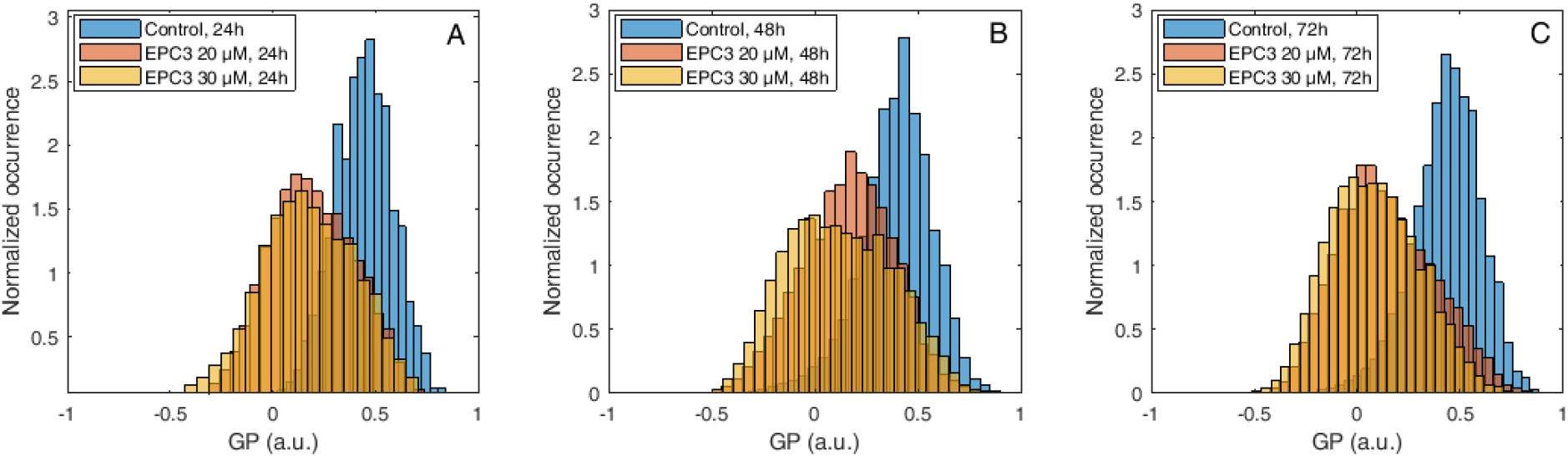
Di-4-ANEPPDQH GP values measured at the PM of MDA-MB-231 cells after EPC3 treatment. Normalized histograms of GP values measured in pixels belonging to the PM of MDA-MB-231 cells labelled with Di-4-ANEPPDQH. Cells were treated with 20 μM (orange bars) or 30 μM (yellow bars) EPC3. GP values measured in cells not treated with EPC3 are shown as blue bars. Fluorescence intensity values were acquired 24 h (Panel A), 48 h (Panel B) and 72 h (Panel C) after the addition of EPC3. For each condition, GP values were pooled from ca. 50 ROIs selected at the PM of distinct cells, in two independent experiments. The total number of calculated GP values (and measured pixels) for each experimental condition was between ca. 10000 and 40000. Measurements were performed at RT. Representative images of treated cells and ROIs are shown in Fig. S1.

As shown in Figs. 3, S2 and Table S1, the fluorescence emission of di-4-ANEPPDQH indicates that EPC3 treatment of cancerous cells induces a time-dependent increase of PM disorder. Only minor differences were observed between IC50 and IC75 EPC3 concentrations (i.e. 20 and 30 μM for MDA-MB-231 cells, respectively). Performing the same experiment using MCF-7 cells (Figs. 4, S3 and Table S2), we observed that EPC3-induced membrane order alterations are stronger and occur faster in MDA-MB-231 cells.

**Figure 4:**
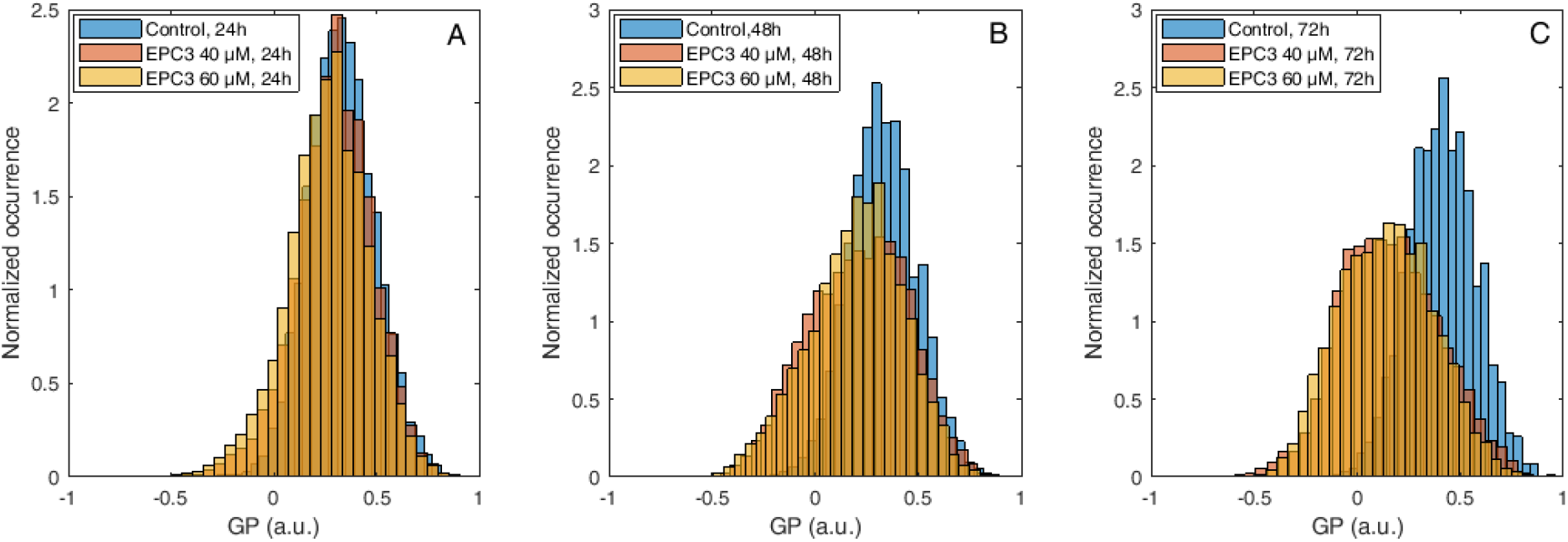
Di-4-ANEPPDQH GP values measured at the PM of MCF-7 cells after EPC3 treatment. Normalized histograms of GP values measured in pixels belonging to the PM of MCF-7 cells labelled with Di-4-ANEPPDQH. Cells were treated with 40 μM (orange bars) or 60 μM (yellow bars) EPC3. GP values measured in cells not treated with EPC3 are shown as blue bars. Fluorescence intensity values were acquired 24 h (Panel A), 48 h (Panel B) and 72 h (Panel C) after the addition of EPC3. For each condition, GP values were pooled from ca. 50 ROIs selected at the PM of distinct cells, in two independent experiments. The total number of calculated GP values (and measured pixels) for each experimental condition was between ca. 10000 and 30000. Measurements were performed at RT. Representative images of treated cells and ROIs are shown in Fig. S1.

In the case of MCF-7 cells, in fact, no significant alteration in membrane order can be observed after 24 h of treatment, both at IC50 and IC75 EPC3 concentrations (i.e. 40 and 60 μM for MCF-7 cells, respectively). After 48h and 72 h however, the GP distribution appears significantly shifted to lower values, independently from EPC3 concentration (Figs. 3 B-C, S3 and Table S2).

Next, we have employed a complementary assay quantifying the GP parameter, following analogous treatment of MDA-MB-231 and MCF-7 cells, using Laurdan as fluorescent probe instead of Di-4-ANEPPDHQ (Fig. S1 I and L).

The results shown in Figs. 5, S4 and Table S3 indicate the presence of a concentration-dependent decrease in membrane order after 48h and 72h treatment with EPC3. We did not observe prominent alterations of the cell membrane order after 24h treatment. These observations are qualitatively similar to those performed using Di-4-ANEPPDQH as fluorescent probe. Also, in analogy to the results shown in Fig. 4 (i.e. labelling of MCF-7 cells with Di-4-ANEPPDQH), the observed changes in lipid packing were larger in MDA-MB-231 than in MCF-7 cells (Fig. S5, S6 and Table S4).

**Figure 5:**
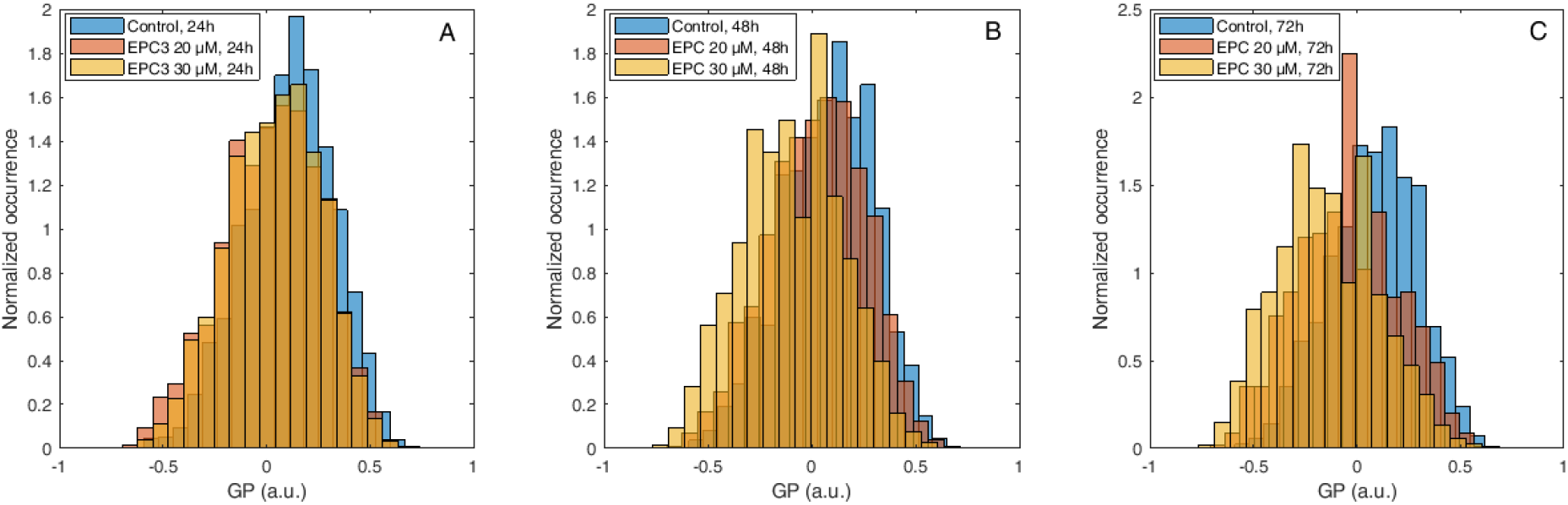
Laurdan GP values measured at the PM of MDA-MB-231 cells after EPC3 treatment. Normalized histograms of GP values measured in pixels belonging to the PM of MDA-MB-231 cells labelled with Laurdan. Cells were treated with 20 μM (orange bars) or 30 μM (yellow bars) EPC3. GP values measured in cells not treated with EPC3 are shown as blue bars. Fluorescence intensity values were acquired 24 h (Panel A), 48 h (Panel B) and 72 h (Panel C) after the addition of EPC3. For each condition, GP values were pooled from ca. 50 ROIs selected at the PM of distinct cells, in two independent experiments. The total number of calculated GP values (and measured pixels) for each experimental condition was between ca. 10000 and 40000. Measurements were performed at RT.

In summary, fluorescence microscopy measurements based on both Laurdan and Di-4-ANEPPDQH labelling indicate a decrease in PM lipid packing and order induced by EPC3, especially in the case of MDA-MB-231-cells. Interestingly, we noticed that the observed effects were much more noticeable when Di-4-ANEPPDHQ was used, instead of Laurdan. This difference might stem from two different factors. First, as previously described (64), Di-4-ANEPPDQH is more effective in specifically labelling the PM. Laurdan, on the other hand, penetrates more easily into the cytosol and labels also intracellular structures (see Fig. S1), thus making ROI selections more difficult and less precise, while generally increasing the spread in observed GP values. Second, Di-4-ANEPPDQH and Laurdan are sensitive to different properties of the lipid bilayer (25). While Laurdan is sensitive to lipid packing, the spectroscopic properties of Di-4-ANEPPDQH are influenced by other factors, such as cholesterol content of the membrane or internal electric dipole potential of the bilayer. It is therefore tempting to assume that the changes in membrane order (observed by using Di-4-ANEPPDQH rather than Laurdan) might be connected to specific alterations in membrane compositions brought about by EPC3. This hypothesis is in agreement with our observation that EPC3 alters the concentration of key lipid components of the PM (e.g. cholesterol and SM) that play an important role in membrane lateral organization (Fig. 2). Similarly, it was shown before that other APLs might influence lipid metabolism and lipid-mediated signaling cascades. On the other hand, our experiments on model membranes clearly indicate that EPC3 can also directly influence lipid-lipid interactions and membrane fluidity, even in the absence of any alteration in membrane composition (Fig. 1).We conclude therefore that EPC3 modulates the physical properties of the PM, by e.g. directly influencing the interactions between cholesterol and other lipids and/or altering the internal electric dipole potential of the bilayer. Additional effects due to alterations in lipid metabolism and PM composition might also be involved *in vivo*.

### EPC3 increases diffusion dynamics of trans-membrane proteins in the PM

We next investigated whether the alterations in lipid-lipid interactions as well as lipid composition caused by EPC3 treatment discussed in the previous paragraphs can have a specific influence on the behavior of membrane proteins. We therefore investigated the diffusive dynamics of two illustrative transmembrane proteins, namely the human EGF receptor (hEGFR) and the Hemagglutinin receptor from Influenza A FP virus (FPV-HA), in MDA-MB-231 cells treated with 30 μM EPC3 (IC75). The hEGFR is a well-characterized membrane protein which partitions in lipid domains of the PM (65). The transmembrane glycoprotein HA of the influenza A virus is known to form trimers at the PM, where it localizes in raft domains (66, 67).

As shown in the previous paragraph, EPC3 (in this case, 30 μM in MDA-MB-231 cells) strongly influences the physical state of the PM, as reported by Di-4-ANEPPDQH. The two examined proteins were labelled with a fluorescent protein (GFP) and visualized via confocal fluorescence microscopy. As shown in Fig. 6 A, EPC3 does not significantly alter the localization of either protein at the PM. We then applied sFCS to quantify the diffusion dynamics of the two fluorescently labelled proteins. Fig. 6 B shows representative autocorrelation curves for the fluorescence signal recorded for hEGFR (upper panel) and FPV-HA (lower panel), also in the presence of EPC3.

**Figure 6:**
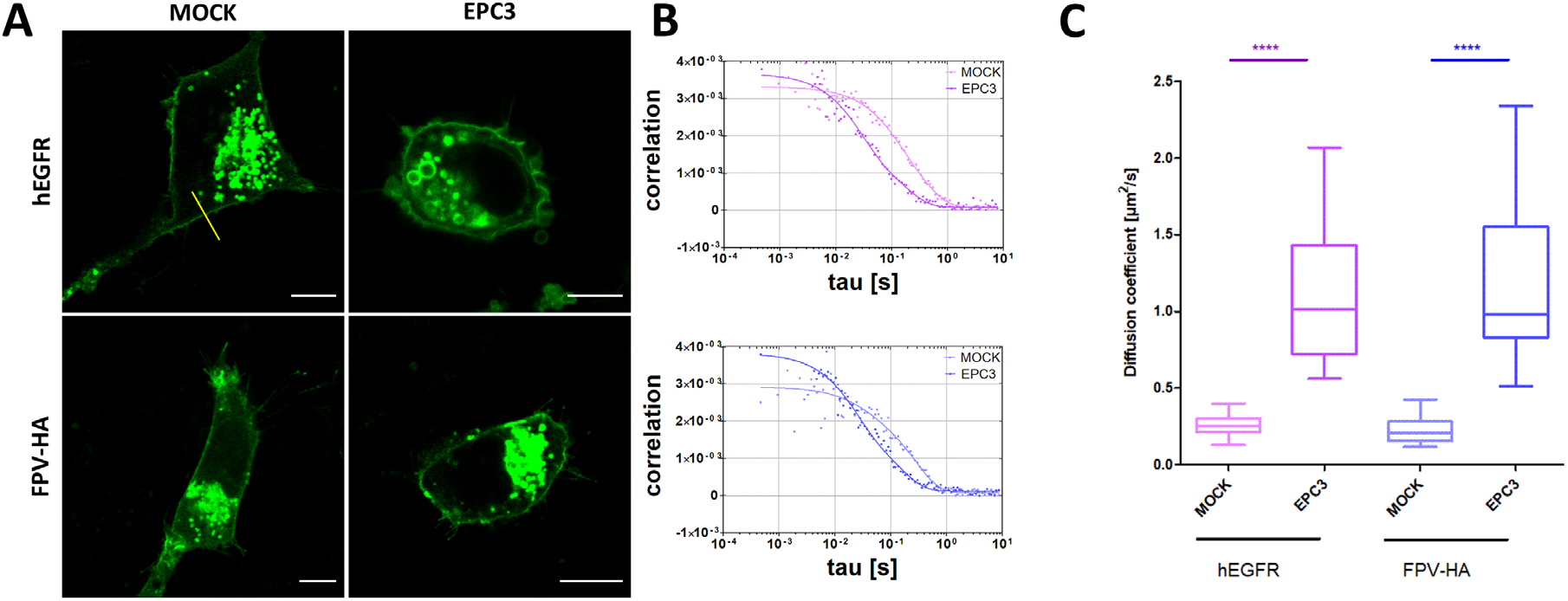
Quantification of protein diffusion in EPC3-treated cells via sFCS. Panel A shows representative confocal fluorescence microscopy images of MDA-MB-231 cells expressing hEGFR-EGFP (upper row), and FPV-HA-EGFP (lower row), in the absence of EPC3 or after a 48 h incubation with EPC3. The yellow line in panel A represents an example of scanning path used for sFCS measurements. Scale bars are 10 μm. Panel B shows representative sFCS autocorrelation functions and fit curves obtained for cells expressing hEGFR-EGFP (upper graph) and FPV-HA-mEGFP (lower graph), in the absence of EPC3 or after a 48 h incubation with EPC3. Fit curves (solid line) were obtained by fitting a two-dimensional diffusion model to the data, as described in the Methods section. Panel C shows box plots of diffusion coefficients calculated from sFCS diffusion times, pooled from three independent experiments (each consisting of measurements on 20 cells). The value of diffusion coefficient D for hEGFR-EGFP significantly increased from 0.26 ± 0.08 μm^2^s^−1^ to 1.1 ± 0.5 μm^2^s^−1^ after EPC3 treatment. The value of diffusion coefficient D for FPV-HA-mEGFP significantly increased from 0.22 ± 0.08 μm^2^s^−1^ to 1.2 ± 0.5 μm^2^s^−1^ after EPC3 treatment. All measurements were performed at RT.

Both membrane proteins diffuse in the PM of untreated cells with a diffusion coefficient D ⁓ 0.25 μm²/s, as expected from previous experiments (19–21). Furthermore, in the presence of EPC3, we observed an average ⁓5-fold increase in diffusion dynamics, both for hEGFR and FPV-HA (Fig. 6 C). Changes in protein diffusion were previously observed as a consequence of alterations in lipid-lipid and lipid-protein interactions. For example, it was shown that hEGFR and HA diffusion coefficients increase as a result of alterations in membrane composition or protein dissociation from membrane domains (28, 65, 68). Our data are compatible with both possibilities. Nevertheless, as already mentioned in the previous paragraph, the results obtained from our experiments in SLBs indicate that changes in membrane composition are not strictly required for the EPC3-induced increase in membrane dynamics.

## CONCLUSIONS

The increased amount of saturated lipids and ordered lipid domains in tumor cells (compared to healthy cells) (69) correlates with a reduction in membrane fluidity/dynamics and an increase in chemotherapy resistance (58, 70). In cancer cells, a wide range of signaling proteins and receptors regulating pro-oncogenic and apoptotic pathways are localized in ordered lipid domains (e.g. raft domains) (6). It is now accepted that APLs exert their effect by interacting with such domains, due to their participation in the regulation of cell survival and cell death pathways (71). In this context, we show for the first time that EPC3 causes alterations in cellular lipid composition and a significant increase in PM disorder. These effects appear stronger in the high-invasive cancer cell line MDA-MB 231 and have a direct effect on the dynamics of membrane components, as demonstrated by our measurements of membrane protein diffusion. Increased protein diffusion dynamics likely affects the likelihood of protein-protein interactions and, therefore, might modulate signaling pathways involved in cell survival. Furthermore, it is possible that the decrease in lipid packing corresponds to a higher permeability of the PM and, thus, an increased intake of EPC3 or other drugs through the cell membrane.

## ACKNOWLEDGMENTS

This work was supported by the grant DN 11/1/2017 from the Bulgarian Science Fund and partially by the grants DО1-154/28/08/2018/ (Scientific Infrastructure on Cell Technologies in Biomedicine (SICTB) from the Bulgarian Ministry of Education and Science) and 254850309 (German Research Foundation (DFG)).

## AUTHOR CONTRIBUTIONS

Conceptualization: R.T., S.C.; Experimental work: R.T., T.S., A.P., D.P., V.U.; Data analysis: R.T.; T.S., A.P., D.P., V.U., S.C.; Writing: R.T., S.C.; Review and Editing: A.P., A.M.; Software: S.C.; Supervision: R.T., S.C.; Funding acquisition: A.M., S.C.

## REFERENCES

1. Jaffres PA, Gajate C, Bouchet AM, Couthon-Gourves H, Chantome A, Potier-Cartereau M, Besson P, Bougnoux P, Mollinedo F, Vandier C. 2016. Alkyl ether lipids, ion channels and lipid raft reorganization in cancer therapy. Pharmacol Ther 165: 114–131.

2. Rios-Marco P, Marco C, Galvez X, Jimenez-Lopez JM, Carrasco MP. 2017. Alkylphospholipids: An update on molecular mechanisms and clinical relevance. Biochim Biophys Acta Biomembr 1859: 1657–1667.

3. Dietrich C, Bagatolli LA, Volovyk ZN, Thompson NL, Levi M, Jacobson K, Gratton E. 2001. Lipid rafts reconstituted in model membranes. Biophys J 80: 1417–1428.

4. Lingwood D, Simons K. 2010. Lipid rafts as a membrane-organizing principle. Science 327: 46–50.

5. Simons K, Toomre D. 2000. Lipid rafts and signal transduction. Nat Rev Mol Cell Biol 1: 31–39.

6. Mollinedo F, Gajate C. 2015. Lipid rafts as major platforms for signaling regulation in cancer. Adv Biol Regul 57: 130–146.

7. Sezgin E, Levental I, Mayor S, Eggeling C. 2017. The mystery of membrane organization: composition, regulation and roles of lipid rafts. Nat Rev Mol Cell Biol 18: 361–374.

8. Mouritsen OG. 2010. The liquid-ordered state comes of age. Biochim Biophys Acta 1798: 1286–1288.

9. Silvius JR. 2005. Partitioning of membrane molecules between raft and non-raft domains: insights from model-membrane studies. Biochim Biophys Acta 1746: 193–202.

10. van der Goot FG, Harder T. 2001. Raft membrane domains: from a liquid-ordered membrane phase to a site of pathogen attack. Semin Immunol 13: 89–97.

11. Ausili A, Martinez-Valera P, Torrecillas A, Gomez-Murcia V, de Godos AM, Corbalan-Garcia S, Teruel JA, Gomez Fernandez JC. 2018. Anticancer Agent Edelfosine Exhibits a High Affinity for Cholesterol and Disorganizes Liquid-Ordered Membrane Structures. Langmuir 34: 8333–8346.

12. Wnetrzak A, Lipiec E, Latka K, Kwiatek W, Dynarowicz-Latka P. 2014. Affinity of alkylphosphocholines to biological membrane of prostate cancer: studies in natural and model systems. J Membr Biol 247: 581–589.

13. Castro BM, Fedorov A, Hornillos V, Delgado J, Acuna AU, Mollinedo F, Prieto M. 2013. Edelfosine and miltefosine effects on lipid raft properties: membrane biophysics in cell death by antitumor lipids. J Phys Chem B 117: 7929–7940.

14. Heczkova B, Slotte JP. 2006. Effect of anti-tumor ether lipids on ordered domains in model membranes. FEBS Lett 580: 2471–2476.

15. Wnetrzak A, Latka K, Makyla-Juzak K, Zemla J, Dynarowicz-Latka P. 2015. The influence of an antitumor lipid - erucylphosphocholine - on artificial lipid raft system modeled as Langmuir monolayer. Mol Membr Biol 32: 189–197.

16. Tanovska M, Rahmani M, Vladimirova Mihaleva L, Berger M, Neshev D, Momchilova A, Tzoneva R. 2019. An Ellipsometric Study of Interaction of Anti-cancer Agent Erufosine on Lipid Model Systems. AIP Conference Proceedings 2075: 170011.

17. Ries J, Chiantia S, Schwille P. 2009. Accurate determination of membrane dynamics with line-scan FCS. Biophys J 96: 1999–2008.

18. Chiantia S, Ries J, Chwastek G, Carrer D, Li Z, Bittman R, Schwille P. 2008. Role of ceramide in membrane protein organization investigated by combined AFM and FCS. Biochim Biophys Acta 1778: 1356–1364.

19. Dunsing V, Mayer M, Liebsch F, Multhaup G, Chiantia S. 2017. Direct evidence of amyloid precursor-like protein 1 trans interactions in cell-cell adhesion platforms investigated via fluorescence fluctuation spectroscopy. Mol Biol Cell 28: 3609–3620.

20. Dunsing V, Chiantia S. 2018. A Fluorescence Fluctuation Spectroscopy Assay of Protein-Protein Interactions at Cell-Cell Contacts. J Vis Exp doi:10.3791/58582.

21. Dunsing V, Luckner M, Zuhlke B, Petazzi RA, Herrmann A, Chiantia S. 2018. Optimal fluorescent protein tags for quantifying protein oligomerization in living cells. Sci Rep 8: 10634.

22. Chiantia S, Ries J, Schwille P. 2009. Fluorescence correlation spectroscopy in membrane structure elucidation. Biochim Biophys Acta 1788: 225–233.

23. Chiantia S, Kahya N, Ries J, Schwille P. 2006. Effects of ceramide on liquid-ordered domains investigated by simultaneous AFM and FCS. Biophys J 90: 4500–4508.

24. Kiessling V, Yang ST, Tamm LK. 2015. Supported lipid bilayers as models for studying membrane domains. Curr Top Membr 75: 1–23.

25. Amaro M, Reina F, Hof M, Eggeling C, Sezgin E. 2017. Laurdan and Di-4-ANEPPDHQ probe different properties of the membrane. J Phys D Appl Phys 50: 134004.

26. Owen DM, Rentero C, Magenau A, Abu-Siniyeh A, Gaus K. 2011. Quantitative imaging of membrane lipid order in cells and organisms. Nat Protoc 7: 24–35.

27. Carter RE, Sorkin A. 1998. Endocytosis of functional epidermal growth factor receptor-green fluorescent protein chimera. J Biol Chem 273:35000–35007.

28. Engel S, Scolari S, Thaa B, Krebs N, Korte T, Herrmann A, Veit M. 2010. FLIM-FRET and FRAP reveal association of influenza virus haemagglutinin with membrane rafts. Biochem J 425: 567–573.

29. Visco I, Chiantia S, Schwille P. 2014. Asymmetric supported lipid bilayer formation via methyl-beta-cyclodextrin mediated lipid exchange: influence of asymmetry on lipid dynamics and phase behavior. Langmuir 30:7475–7484.

30. Hofer CT, Di Lella S, Dahmani I, Jungnick N, Bordag N, Bobone S, Huang Q, Keller S, Herrmann A, Chiantia S. 2019. Structural determinants of the interaction between influenza A virus matrix protein M1 and lipid membranes. Biochim Biophys Acta Biomembr 1861: 1123–1134.

31. Stoyanova T, Uzunova V, Popova D, Hadzhilazova M, Berger MR, Momchilova A, Toshkova R, Tzoneva R. 2018. Effect of Erufosine on MDA-MB 231 Breast Cancer Cells. Journal of Oncology Research Forecast 1: 105.

32. Pankov R, Markovska T, Antonov P, Ivanova L, Momchilova A. 2006. The plasma membrane lipid composition affects fusion between cells and model membranes. Chem Biol Interact 164: 167–173.

33. Bligh EG, Dyer WJ. 1959. A rapid method of total lipid extraction and purification. Can J Biochem Physiol 37:911–917.

34. Kahovcova J, Odavic R. 1969. A simple method for the quantitative analysis of phospholipids separated by thin layer chromatography. J Chromatogr 40: 90–96.

35. Nikolova-Karakashian MN, Petkova H, Koumanov KS. 1992. Influence of cholesterol on sphingomyelin metabolism and hemileaflet fluidity of rat liver plasma membranes. Biochimie 74: 153–159.

36. Petrasek Z, Schwille P. 2008. Precise measurement of diffusion coefficients using scanning fluorescence correlation spectroscopy. Biophys J 94:1437–1448.

37. Ries J, Schwille P. 2006. Studying slow membrane dynamics with continuous wave scanning fluorescence correlation spectroscopy. Biophys J 91: 1915–1924.

38. Dorlich RM, Chen Q, Niklas Hedde P, Schuster V, Hippler M, Wesslowski J, Davidson G, Nienhaus GU. 2015. Dual-color dual-focus line-scanning FCS for quantitative analysis of receptor-ligand interactions in living specimens. Sci Rep 5: 10149.

39. Chiantia S, Ries J, Kahya N, Schwille P. 2006. Combined AFM and two-focus SFCS study of raft-exhibiting model membranes. Chemphyschem 7: 2409–2418.

40. Veatch SL, Keller SL. 2005. Seeing spots: complex phase behavior in simple membranes. Biochim Biophys Acta 1746: 172–185.

41. Sengupta P, Baird B, Holowka D. 2007. Lipid rafts, fluid/fluid phase separation, and their relevance to plasma membrane structure and function. Semin Cell Dev Biol 18:583–590.

42. Gomide AB, Thome CH, dos Santos GA, Ferreira GA, Faca VM, Rego EM, Greene LJ, Stabeli RG, Ciancaglini P, Itri R. 2013. Disrupting membrane raft domains by alkylphospholipids. Biochim Biophys Acta 1828: 1384–1389.

43. Boggs K, Rock CO, Jackowski S. 1998. The antiproliferative effect of hexadecylphosphocholine toward HL60 cells is prevented by exogenous lysophosphatidylcholine. Biochim Biophys Acta 1389:1–12.

44. Jimenez-Lopez JM, Rios-Marco P, Marco C, Segovia JL, Carrasco MP. 2010. Alterations in the homeostasis of phospholipids and cholesterol by antitumor alkylphospholipids. Lipids Health Dis 9: 33.

45. Wieder T, Orfanos CE, Geilen CC. 1998. Induction of ceramide-mediated apoptosis by the anticancer phospholipid analog, hexadecylphosphocholine. J Biol Chem 273: 11025–11031.

46. Geilen CC, Wieder T, Reutter W. 1992. Hexadecylphosphocholine inhibits translocation of CTP:choline-phosphate cytidylyltransferase in Madin-Darby canine kidney cells. J Biol Chem 267:6719–6724.

47. Marco C, Jimenez-Lopez JM, Rios-Marco P, Segovia JL, Carrasco MP. 2009. Hexadecylphosphocholine alters nonvesicular cholesterol traffic from the plasma membrane to the endoplasmic reticulum and inhibits the synthesis of sphingomyelin in HepG2 cells. Int J Biochem Cell Biol 41: 1296–1303.

48. Colombini M. 2013. Membrane channels formed by ceramide. Handb Exp Pharmacol doi:10.1007/978-3-7091-1368-4_6:109-126.

49. Birge RB, Boeltz S, Kumar S, Carlson J, Wanderley J, Calianese D, Barcinski M, Brekken RA, Huang X, Hutchins JT, Freimark B, Empig C, Mercer J, Schroit AJ, Schett G, Herrmann M. 2016. Phosphatidylserine is a global immunosuppressive signal in efferocytosis, infectious disease, and cancer. Cell Death Differ 23: 962–978.

50. Danilo C, Gutierrez-Pajares JL, Mainieri MA, Mercier I, Lisanti MP, Frank PG. 2013. Scavenger receptor class B type I regulates cellular cholesterol metabolism and cell signaling associated with breast cancer development. Breast Cancer Res 15: R87.

51. Antalis CJ, Uchida A, Buhman KK, Siddiqui RA. 2011. Migration of MDA-MB-231 breast cancer cells depends on the availability of exogenous lipids and cholesterol esterification. Clin Exp Metastasis 28: 733–741.

52. Awad AB, Fink CS, Williams H, Kim U. 2001. In vitro and in vivo (SCID mice) effects of phytosterols on the growth and dissemination of human prostate cancer PC-3 cells. Eur J Cancer Prev 10: 507–513.

53. Carrasco MP, Jimenez-Lopez JM, Rios-Marco P, Segovia JL, Marco C. 2010. Disruption of cellular cholesterol transport and homeostasis as a novel mechanism of action of membrane-targeted alkylphospholipid analogues. Br J Pharmacol 160: 355–366.

54. Ait Slimane T, Hoekstra D. 2002. Sphingolipid trafficking and protein sorting in epithelial cells. FEBS Lett 529: 54–59.

55. Abramczyk H, Surmacki J, Kopec M, Olejnik AK, Lubecka-Pietruszewska K, Fabianowska-Majewska K. 2015. The role of lipid droplets and adipocytes in cancer. Raman imaging of cell cultures: MCF10A, MCF7, and MDA-MB-231 compared to adipocytes in cancerous human breast tissue. Analyst 140: 2224–2235.

56. de Gonzalo-Calvo D, Lopez-Vilaro L, Nasarre L, Perez-Olabarria M, Vazquez T, Escuin D, Badimon L, Barnadas A, Lerma E, Llorente-Cortes V. 2015. Intratumor cholesteryl ester accumulation is associated with human breast cancer proliferation and aggressive potential: a molecular and clinicopathological study. BMC Cancer 15: 460.

57. Bozza PT, Viola JP. 2010. Lipid droplets in inflammation and cancer. Prostaglandins Leukot Essent Fatty Acids 82: 243–250.

58. Rysman E, Brusselmans K, Scheys K, Timmermans L, Derua R, Munck S, Van Veldhoven PP, Waltregny D, Daniels VW, Machiels J, Vanderhoydonc F, Smans K, Waelkens E, Verhoeven G, Swinnen JV. 2010. De novo lipogenesis protects cancer cells from free radicals and chemotherapeutics by promoting membrane lipid saturation. Cancer Res 70: 8117–8126.

59. Wang RF, Zhang LH, Shan LH, Sun WG, Chai CC, Wu HM, Ibla JC, Wang LF, Liu JR. 2013. Effects of the fibroblast activation protein on the invasion and migration of gastric cancer. Exp Mol Pathol 95: 350–356.

60. Li YC, Park MJ, Ye SK, Kim CW, Kim YN. 2006. Elevated levels of cholesterol-rich lipid rafts in cancer cells are correlated with apoptosis sensitivity induced by cholesterol-depleting agents. Am J Pathol 168: 1107–1118; quiz 1404-1105.

61. Petit K, Suwalsky M, Colina JR, Aguilar LF, Jemiola-Rzeminska M, Strzalka K. 2019. In vitro effects of the antitumor drug miltefosine on human erythrocytes and molecular models of its membrane. Biochim Biophys Acta Biomembr 1861: 17–25.

62. Pehlivanova V, Uzunova V, Tsoneva I, Berger MR, Ugrinova I, Tzoneva R. 2013. Effect of Erufosine on the Reorganization of Cytoskeleton and Cell Death in Adherent Tumor and Non-Tumorigenic Cells. Biotechnology & Biotechnological Equipment 27: 3695–3699.

63. Stoyanova T, Uzunova V, Momchilova A, Tzoneva R, Ugrinova I. 2018. The treatment of breast cancer cells with erufosine leads to actin cytoskeleton reorganization, inhibition of cell motility, cell cycle arrest and apoptosis. Compt rend Acad Bulg Sci.

64. Mazeres S, Fereidouni F, Joly E. 2017. Using spectral decomposition of the signals from laurdan-derived probes to evaluate the physical state of membranes in live cells. F1000Res 6:763.

65. Bag N, Huang S, Wohland T. 2015. Plasma Membrane Organization of Epidermal Growth Factor Receptor in Resting and Ligand-Bound States. Biophys J 109: 1925–1936.

66. Takeda M, Leser GP, Russell CJ, Lamb RA. 2003. Influenza virus hemagglutinin concentrates in lipid raft microdomains for efficient viral fusion. Proc Natl Acad Sci U S A 100: 14610–14617.

67. Scheiffele P, Rietveld A, Wilk T, Simons K. 1999. Influenza viruses select ordered lipid domains during budding from the plasma membrane. J Biol Chem 274: 2038–2044.

68. Shvartsman DE, Kotler M, Tall RD, Roth MG, Henis YI. 2003. Differently anchored influenza hemagglutinin mutants display distinct interaction dynamics with mutual rafts. J Cell Biol 163: 879–888.

69. Beloribi-Djefaflia S, Vasseur S, Guillaumond F. 2016. Lipid metabolic reprogramming in cancer cells. Oncogenesis 5: e189.

70. Ollila S, Hyvonen MT, Vattulainen I. 2007. Polyunsaturation in lipid membranes: dynamic properties and lateral pressure profiles. J Phys Chem B 111: 3139–3150.

71. Kostadinova A, Topouzova-Hristova T, Momchilova A, Tzoneva R, Berger MR. 2015. Antitumor Lipids--Structure, Functions, and Medical Applications. Adv Protein Chem Struct Biol 101: 27–66.

