## Supplementary Info for "Effect of Erufosine on Membrane Lipid Order in Breast Cancer Cell Models"

### SUPPLEMENTARY INFORMATION

**Tzoneva R., Stoyanova T. et al.**

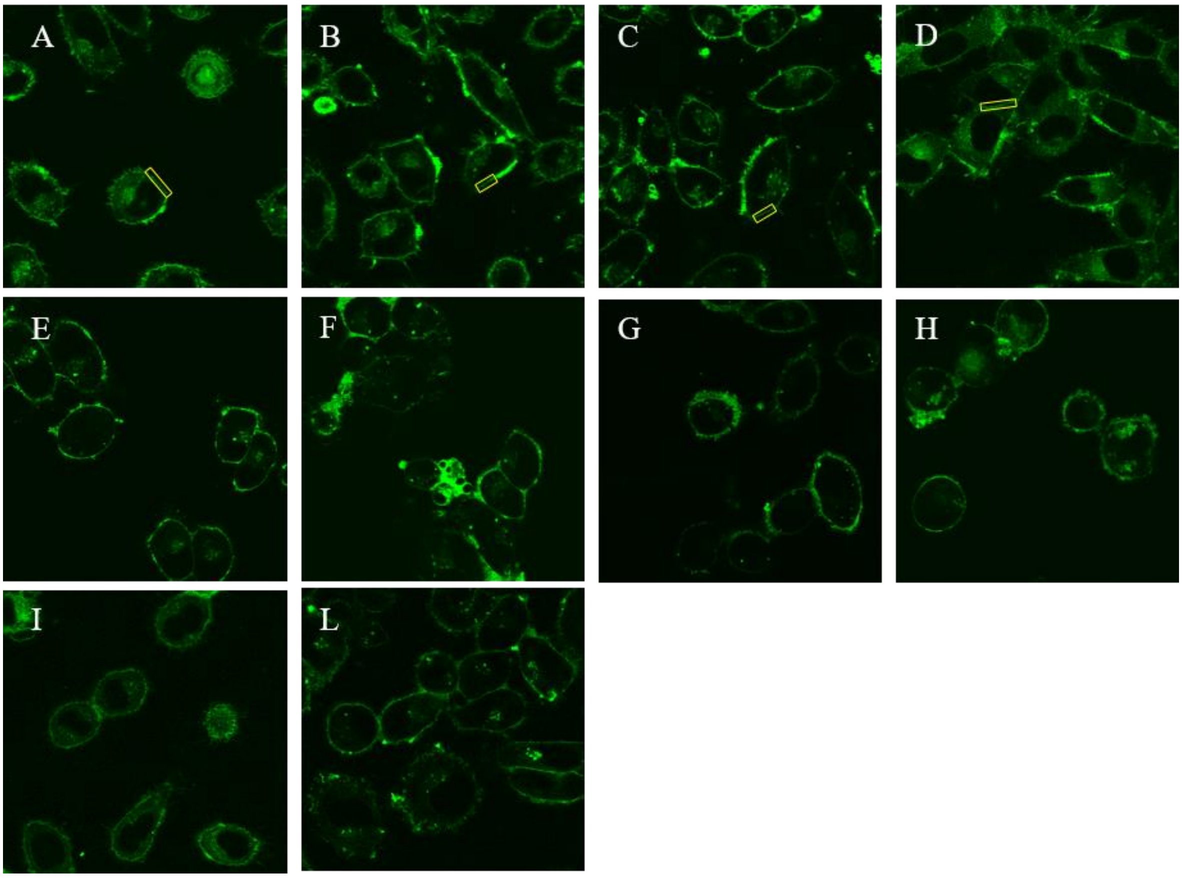


**Figure S1: Typical fluorescence microscopy images of cells samples used in this work.** Confocal microscopy images of MDA-MB 231 (panels A-D, I) and MCF-7 cells (panels E-H, L). Cells were labelled with Di-4-ANEPPDHQ (panels A-H) or with Laurdan (panels I and L). Samples are shown before treatment (panels A,E, I and L) and after treatment with EPC3 (24h: panels B and F; 48h: panels C and G; 72h: panels D and H. The yellow rectangles in panels A-D represent examples of ROIs selected for general polarization analysis. All images are 106 µm x 106 µm.

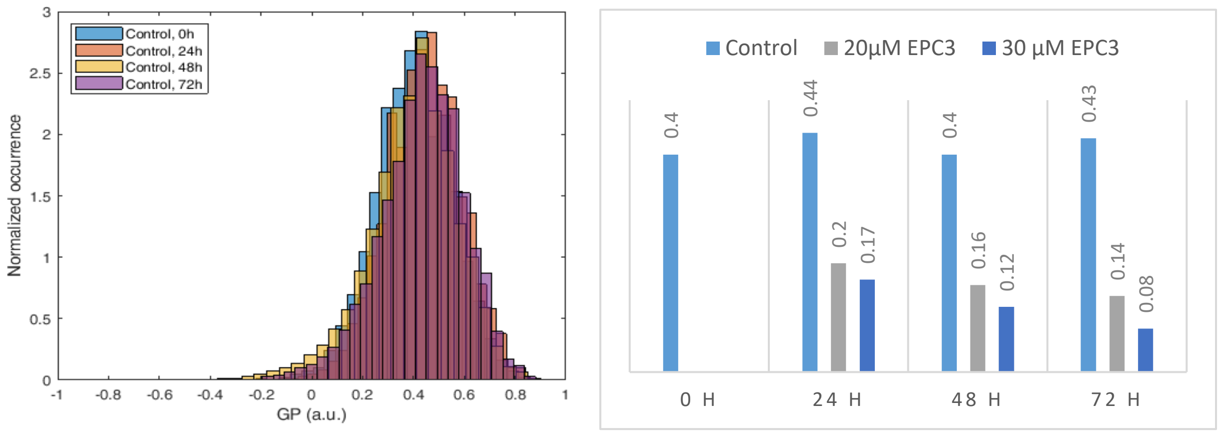


**Figure S2: Analysis of Di-4-ANEPPDHQ GP values measured at the PM of MDA-MB-231.** Left panel: GP values for all the control samples at different time points (see Fig. 2 in the main text), including the data acquired before the beginning of the EPC3 treatment (Control, 0h). Right panel: Average values of the normalized histograms shown in Fig. 2 and in Table S1, as a function of treatment time.

| | Mean | Std. dev. |
|---|---|---|
| Control 0h | 0.40 | 0.15 |
| Control 24h | 0.44 | 0.15 |
| IC50 24h | 0.20 | 0.22 |
| IC75 24h | 0.17 | 0.24 |
| Control 48h | 0.40 | 0.17 |
| IC50 48h | 0.16 | 0.22 |
| IC75 48h | 0.12 | 0.26 |
| Control 72h | 0.43 | 0.17 |
| IC50 72h | 0.14 | 0.23 |
| IC75 72h | 0.08 | 0.22 |

**Table S1**

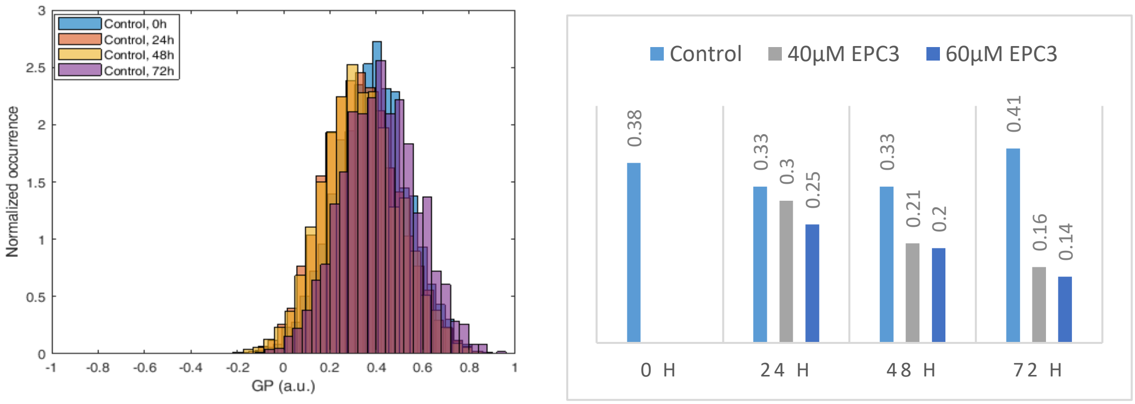


**Figure S3: Analysis of Di-4-ANEPPDHQ GP values measured at the PM of MCF-7.** Left panel: GP values for all the control samples at different time points (see Fig. 3 in the main text), including the data acquired before the beginning of the EPC3 treatment (Control, 0h). Right panel: Average values of the normalized histograms shown in Fig. 3 and in Table S2, as a function of treatment time.

| | Mean | Std. dev. |
|---|---|---|
| Control 0h | **0.38** | **0.16** |
| Control 24h | **0.33** | **0.16** |
| IC50 24h | **0.30** | **0.19** |
| IC75 24h | **0.25** | **0.20** |
| Control 48h | **0.33** | **0.16** |
| IC50 48h | **0.21** | **0.24** |
| IC75 48h | **0.20** | **0.23** |
| Control 72h | **0.41** | **0.16** |
| IC50 72h | **0.16** | **0.24** |
| IC75 72h | **0.14** | **0.23** |

**Table S2**

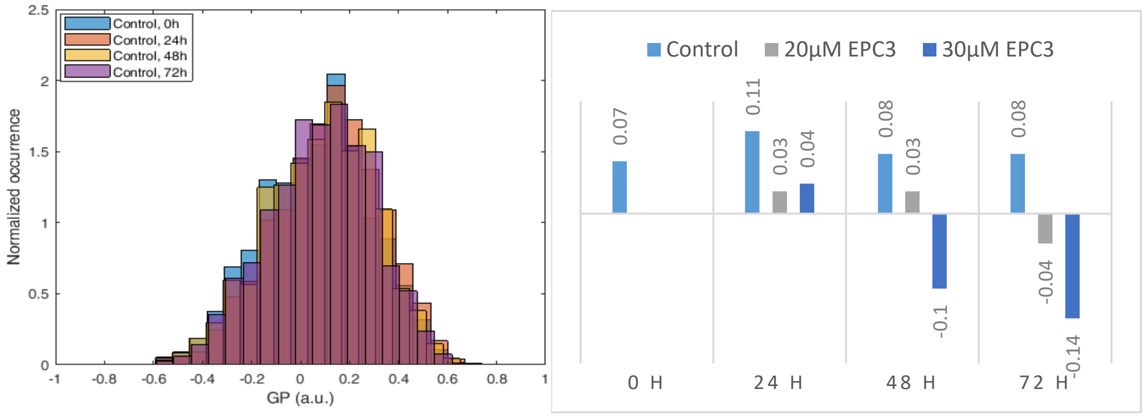


**Figure S4: Analysis of Laurdan GP values measured at the PM of MDA-MB-231.** Left panel: GP values for all the control samples at different time points (see Fig. 4 in the main text), including the data acquired before the beginning of the EPC3 treatment (Control, 0h). Right panel: Average values of the normalized histograms shown in Fig. 4 and in Table S3, as a function of treatment time.

| | Mean | Std. dev. |
|---|---|---|
| Control 0h | **0.07** | **0.22** |
| Control 24h | **0.11** | **0.22** |
| IC50 24h | **0.03** | **0.24** |
| IC75 24h | **0.04** | **0.23** |
| Control 48h | **0.08** | **0.22** |
| IC50 48h | **0.03** | **0.23** |
| IC75 48h | **-0.10** | **0.24** |
| Control 72h | **0.08** | **0.21** |
| IC50 72h | **-0.04** | **0.24** |
| IC75 72h | **-0.14** | **0.24** |

**Table S3**

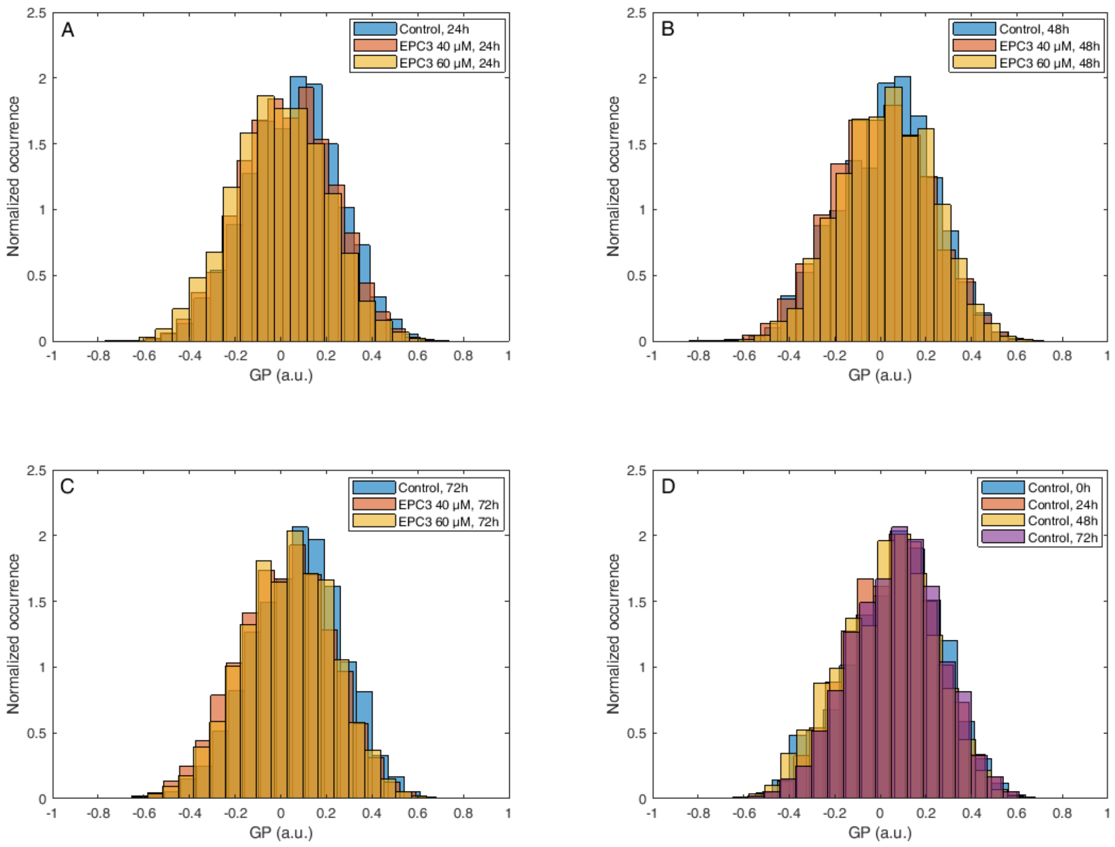


**Figure S5: Laurdan GP values measured at the PM of MCF-7 cells after EPC3 treatment.** Panels A-C: normalized histograms of GP values measured in pixels belonging to the PM of MCF-7 cells labelled with Laurdan. Cells were treated with 40 µM (orange bars) or 60 µM (yellow bars) EPC3. GP values measured in cells not treated with EPC3 are shown as blue bars. Fluorescence intensity values were acquired 24 h (Panel A), 48 h (Panel B) and 72 h (Panel C) after the addition of EPC3. Panel D shows GP values for all the control samples at different time points, including the data acquired before the beginning of the EPC3 treatment (Control, 0h). For each condition, GP values were pooled from ca. 50 ROIs selected at the PM of distinct cells, in two independent experiments. The total number of calculated GP values (and measured pixels) for each experimental condition was between ca. 10000 and 30000. Measurements were performed at RT.

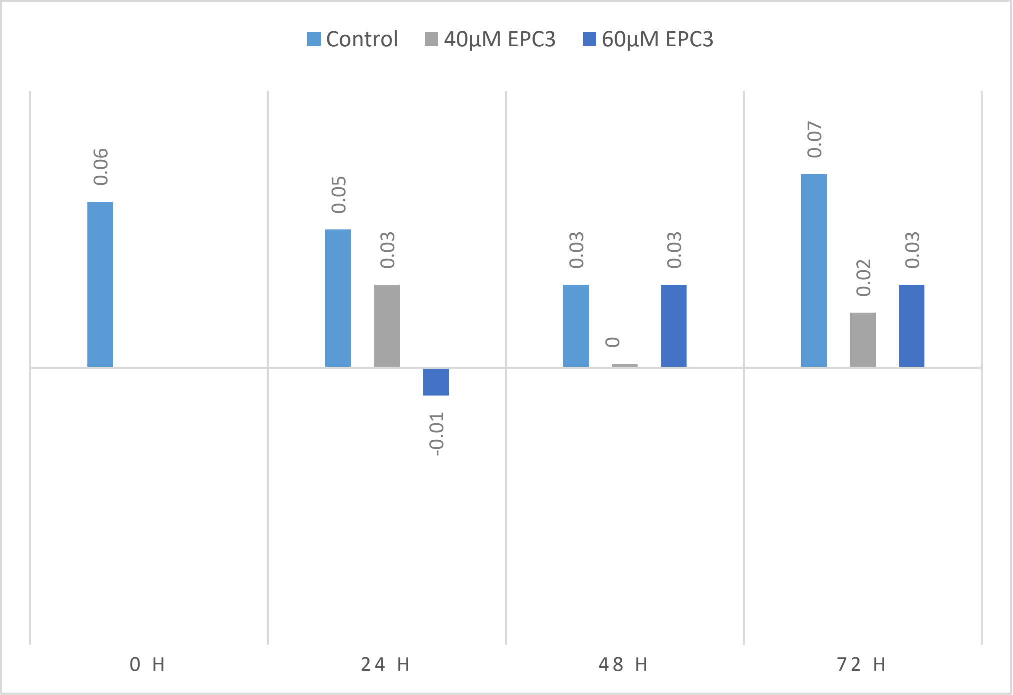


**Figure S6: Analysis of Laurdan GP values measured at the PM of MCF-7 cells.** Average values of the normalized histograms shown in Fig. S5 and in Table S4, as a function of treatment time.

| | Mean | Std. dev. |
|---|---|---|
| Control 0h | **0.06** | **0.21** |
| Control 24h | **0.05** | **0.20** |
| IC50 24h | **0.03** | **0.20** |
| IC75 24h | **-0.01** | **0.21** |
| Control 48h | **0.03** | **0.21** |
| IC50 48h | **0.00** | **0.21** |
| IC75 48h | **0.03** | **0.20** |
| Control 72h | **0.07** | **0.20** |
| IC50 72h | **0.02** | **0.21** |
| IC75 72h | **0.03** | **0.20** |

**Table S4**
